# Zebrafish model of *tcn2* deletion reveals new molecular insights into the role of vitamin B12 in embryonic development

**DOI:** 10.1101/2023.10.14.562325

**Authors:** Ajay Deepak Verma, Suraj S Nongmaithem, Challapalli Mounika, Swetha Ramachandran, Anushri Umesh, Giriraj Ratan Chandak

## Abstract

Vitamin B12 (B12) is one of the key co-factors in One-Carbon Metabolism pathway, and its dysregulation is associated with various clinical conditions including congenital malformations, neural tube defects, and low birthweight, etc. However, the underlying molecular mechanism is scarcely studied. This study investigated the role of B12 in early embryonic development by generating a knockout of transcobalamin 2 (*tcn2*), a B12 transporter in zebrafish.

We generated *tcn2*^-/-^ zebrafish by creating premature stop codons in exon 7 and confirmed by qRT-PCR and immunoblotting. Phenotypic changes were captured, growth and survival assays were conducted at different developmental stages. B12 supplementation assay was conducted by rearing *tcn2*^-/-^ embryos in embryo-medium containing 10 µM cyanocobalamin. RNA-sequencing data was generated in triplicates in embryos at 1-cell and 24 hpf stages, with and without B12 supplementation. Genes with log_2_FC ≥ 0.6 and adjusted p-value ≤ 0.01 were considered differentially expressed genes (DEGs) and proceeded for functional annotation and enrichment analysis.

The F1 *tcn2^-/-^* embryos grew normally. However, the F2 *tcn2^-/-^* embryos remained unhatched, showed delayed somitogenesis, curved or deformed tails, abnormal yolk extension, pericardial and yolk sac edema, etc. and ∼70% were dead by 4 dpf. The abnormalities were partially ameliorated by B12 supplementation throughout developmental stages. Transcriptome analysis identified DEGs enriched in pathways including spliceosome, ribosome biogenesis, branched-chain amino acid degradation, lysosome, iron homeostasis, metabolic and immune response pathways, etc. We observed dysregulated expression of many genes with epigenetic functions including *sfswap*, *ctcf*, *mettl3*, *alkbh5*, and *hdac*, *etc.,* in the 1-cell *tcn2*^-/-^ embryos. Notably, B12 supplementation restored expression of many DEGs indicating a potential role of B12 in transcriptional regulation during embryonic development.

Our study delineates the importance of B12 during embryonic development and provides interesting insights into the molecular mechanisms underlying the observed phenotypic changes.

## Introduction

Vitamin B12 (B12) is one of the key co-factors in two essential pathways methyl malonyl-CoA synthesis, and One-Carbon Metabolism (OCM) pathway which influences various biological processes including DNA synthesis, amino acid homeostasis, epigenetic regulation, and redox defense. It is an essential micronutrient and individuals with B12 deficiency or having mutations in genes involved in its absorption, transport, and/or intracellular processing, etc., develop congenital defects, megaloblastic anaemia, neurological and psychological dysfunctions, etc.[1]. In humans, B12 is present in two biologically active forms i.e., 5’-deoxyadenosylcobalamin and methylcobalamin [2]. 5’-deoxyadenosylcobalamin is a cofactor for the mitochondrial enzyme methylmalonyl-CoA mutase which catalyzes the conversion of methylmalonyl-coenzyme A to succinyl-coA [2] an intermediary of the citric acid cycle [3]. Methylcobalamin acts as a cofactor for the cytosolic enzyme methionine synthase which catalyzes the conversion of homocysteine to methionine and N5-methyl tetrahydrofolate to tetrahydrofolate thus plays a crucial role in regulating the OCM pathway [4]. Elevated homocysteine levels, an established marker of disturbed OCM pathway, are associated with neural tube defects, cognitive impairment of the child, congenital heart defects, and many other diseases [5]. We and others have shown that a low maternal B12 status, especially during pregnancy is associated with low birth weight and fetal growth restriction, increased risk of neural tube defects, neurocognitive developmental deficits, and increased insulin resistance in their offspring [6–8].

Synthesis of B12 is limited to a few prokaryotes therefore humans completely depend on the food intake and gut microbiota. Thus, nutritional, environmental, and genetic factors are the key determinants of B12 levels. Genetic association studies in humans have identified many genetic variants in B12 transporters (*TCN1, TCN2, IF*) and other genes including *FUT2, FUT6, CUBN, CD320, MUT,* and *MMAA* etc., to be associated with plasma B12 levels [9,10]. Haptocorrin encoded by *TCN1*, intrinsic factor by *IF*, and transcobalamin II by *TCN2* are three paralogous proteins involved in B12 assimilation, absorption, and transport in human beings. In zebrafish, transcobalamin 2 (*tcn2*) is the only known B12 transporter and has a similar structure and B12-binding affinity with human *TCN2*. It shares ∼30% sequence identity at the protein level with all three human paralogs [11]. Zebrafish have been used as a tractable model to study embryonic development as they produce large clutches of *ex-utero* developing optically transparent embryos. We used zebrafish as a model organism to study the effect of B12 deficiency in fetal developmental programming.

In this study, we generated a B12-deficient zebrafish model by CRISPR/Cas9 mediated elimination of *tcn2* expression and investigated phenotypic changes through generations. Further, we also attempted to understand underlying gene expression changes and the effect of B12 supplementation on the phenotype and gene expression changes.

## Materials and Methods

Adult zebrafish (Tubingen strain) were raised and maintained in a temperature controlled (28±1°C) room with a 14:10 light/dark cycle as described earlier [12]. The embryos were staged by hours post fertilization (hpf) and by morphological criteria as described [13]. All zebrafish experiments were conducted as per Institutional guidelines and approved by institutional ethics committee.

### Generation of *tcn2*^-/-^ zebrafish lines

CRISPR Cas9 genome editing was performed as described earlier [14]. Guide RNAs (gRNAs) showing minimal off-target sites were designed using CHOPCHOP (http://chopchop.cbu.uib.no). Oligos encoding gRNAs targeting exon 7 of *tcn2* were synthesized by M/s. Bioserve India and were cloned into DR274 plasmid (Addgene, USA) (Supplementary Table 1). After confirmation of the sequence of clones by Sanger sequencing, gRNAs were synthesized using MAXIscript T7 Transcription Kit (Invitrogen, USA) following the manufacturer’s protocol. One-cell stage embryos were injected with ∼1 nl solution containing 100 pg of gRNA and 200 pg of GeneArt Platinum Cas9 protein (Thermo Fisher Scientific, USA). Injected embryos were raised into adults, genotyped, and backcrossed to identify transmitted mutations in the G1 progeny. Homozygotes were generated by breeding G2 heterozygotes (Figure 1). The primer sequences used for genotyping by Sanger sequencing are listed in Supplementary Table 1.

**Figure 1:**
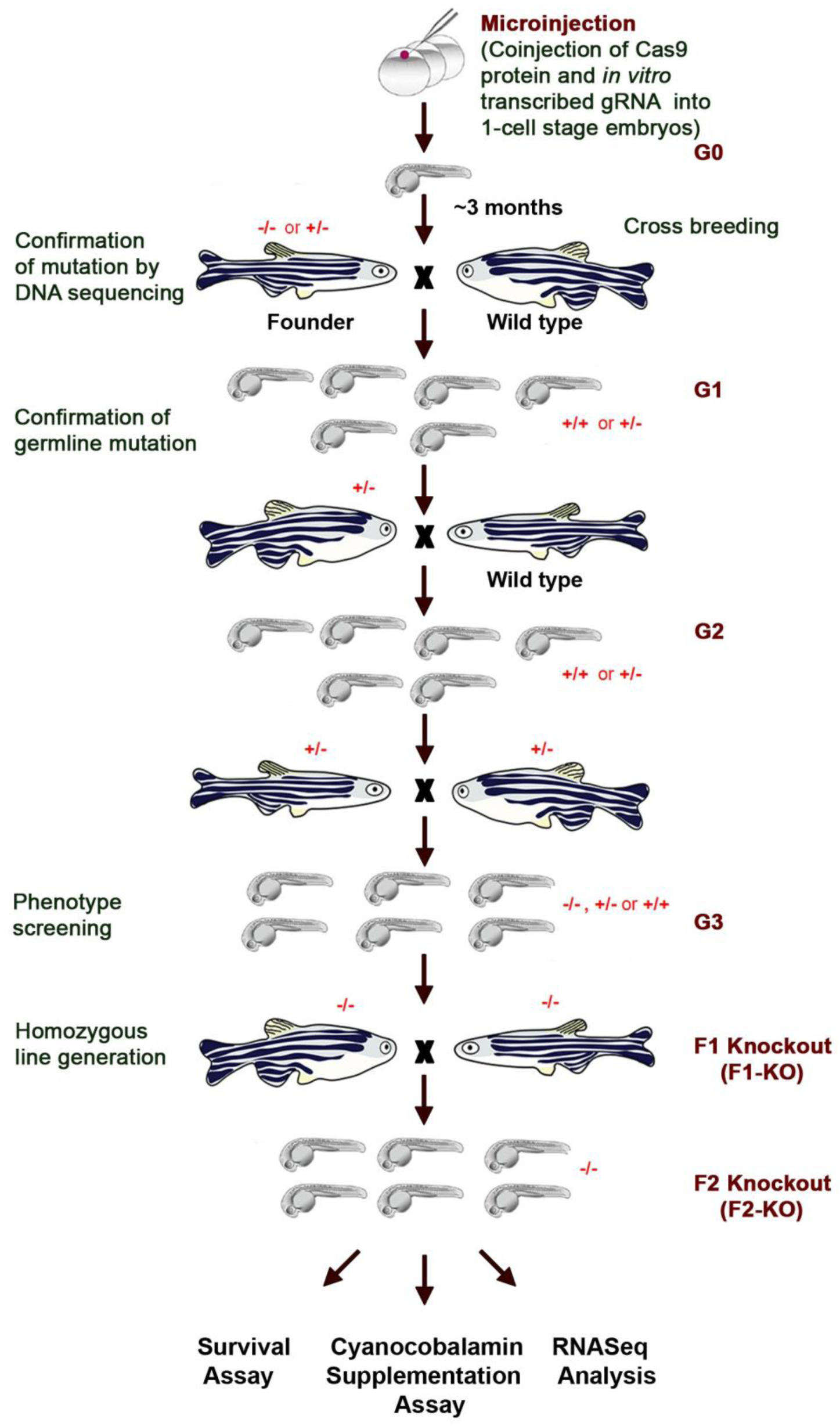
Study design. – Generation of *tcn2*^-/-^ zebrafish using CRISPR/Cas9

### B12 supplementation and imaging of zebrafish embryos

The embryos were collected within 30 min of the start of breeding and transferred to the Petri dishes (80 mm) containing embryo medium (E3) at a density of 50 embryos/plate. A commercially available form of vitamin B12, Cyanocobalamin (10 µM) was added to the E3 medium as needed. All Petri dishes were kept in dark, and the dead embryos were scored. A total of 1594 *tcn2*^-/-^ embryos (700 untreated and 894 treated with B12) were analyzed in four separate experimental sets. The embryos were dechorionated manually and anesthetized in embryo water containing 0.17 mg/ml tricaine. The brightfield images were captured using a stereo microscope Leica M205A (Leica Microsystems GmbH, Germany).

### RNA isolation, RNA-Sequencing and transcriptome analysis

Total RNA was isolated from zebrafish embryos using TRIzol reagent (Invitrogen, USA) following the manufacturer’s instructions. After DNase treatment, RNA was purified by a phenol/chloroform extraction step. The quality of RNA was checked by gel electrophoresis and quantitation done by NanoDrop 2000 (Thermo Scientific, USA). For the RNA sequencing experiments, the RNA quality was assessed using 4200 TapeStation (Agilent, USA).

A total of 15 RNA samples from zebrafish embryos – biological triplicates of 1-cell stage wild-type (Wt) and *tcn2*^-/-^ (KO); and 24 hpf stage Wt, KO and B12 supplemented KO (KO+B12) embryos were subjected to RNA-sequencing analysis. Briefly, 1µg of total RNA (RIN > 8.0) was enriched using Poly(A) RNA Selection kit v1.5 and RNA libraries were prepared using CORALL mRNA-Seq Library Prep kit (Lexogen GmbH, Austria) according to manufacturer’s protocols. After purification, the library was amplified using specific oligos (Lexogen GmbH, Austria) required for cluster generation to create the final cDNA library. Libraries were sequenced as paired-end 150 bp reads on NovaSeq 6000 (Illumina, USA) following prescribed protocols.

Data analysis was performed using European Galaxy Server. In brief, FASTQ.GZ files were processed in FastP (Version 0.23.2) [15] using default set parameters except for qualified quality Phred Q30, and a minimum read length filter of 20 bases (Supplementary Table S2A). For each sample, about 40 million reads were mapped uniquely (Supplementary Table S2B). The processed paired-end reads were aligned to the zebrafish genome (GRCz11.107.chr.gtf) by RNAstar (Version 2.7.8a) [16]. The BAM file generated were then assembled into the transcriptome (GTF file) using StringTie (Version 2.1.7) [17]. From the gene counts output table, differential gene expression was estimated using DESeq2 based on a model using the negative binomial distribution [18]. Genes with an adjusted p-value ≤ 0.01 (Benjamini and Hochberg method); log_2_FC ≥ 0.6 were considered differentially expressed. Functional annotation and enrichment analyses were performed on differentially expressed genes (DEGs) using DAVID [19].

### Quantitative Real-time PCR

Using Superscript II (Invitrogen, USA), 3 µg of total RNA isolated from zebrafish embryos was reverse transcribed at 42^°^C for 50 min following the manufacturer’s instructions. Quantitative PCR was carried out using the following conditions: 50^°^C: 2 min, 95^°^C: 10 min, and 40 cycles of 95^°^C: 15s, 60^°^C: 25s, and 72^°^C: 30s. For each sample, dissociation step was performed at 95^°^C: 15s, 60^°^C: 1 min, and 95^°^C: 15s at the end of the amplification phase (Supplementary Table 1). Amplification of PCR products was quantified using power SYBR green dye (Applied Biosystems, USA) and fluorescence was monitored on ABI ViiA 7 and QuantStudio 5 Realtime PCR system (Applied Biosystems, USA). Melting curve analysis was done for each amplicon. The 2^-ΔΔCt^ method was used for quantitation with *elf1a* and *tubulin1a* as housekeeping genes. Ct values for all housekeeping genes were consistent for all the samples. The experiments for each gene were done in triplicate and analysis of three independent biological replicates was performed.

### Immunoblotting

Zebrafish embryos or tissues were crushed and homogenized in Laemmli sample buffer and boiled for 10 min. The protein lysates were quantified by Amido Black staining and ∼30 µg lysates were separated on 10% SDS-polyacrylamide gels and electroblotted onto the polyvinylidene difluoride (PVDF) membrane. Upon transfer, the PVDF membrane was blocked with 5% skimmed milk in Tris-Buffered Saline containing 0.1% Tween-20 (TBST) for 1 hr at room temperature. The blot was incubated with a primary antibody diluted in TBST. After the washing steps, the blot was incubated with the appropriate secondary antibody for 1 hr and visualized on a chemiluminescence system using ECL chemiluminescent substrate reagent kit (Thermo Fisher Scientific, USA). A rabbit polyclonal antibody was raised in-house against His-tagged zebrafish tcn2 protein excluding the signal peptide. Immunoblot analysis using this antibody allowed the detection of tcn2 in different embryonic stages and adult tissues.

## Results

We generated a zebrafish model of B12 deficiency by generating knockout of *tcn2* using CRISPR/Cas9 method and tried to understand the molecular changes at transcriptome levels of the knockout phenotypes at two different embryonic stages to understand the role of B12 in embryonic development.

### Generation and characterization of *tcn2*^-/-^ zebrafish line

Study design and generation of *tcn2*^-/-^ is shown in Figure 1. We generated two loss-of-function alleles of *tcn2*^-/-^ (d13 and d20 lines) in zebrafish using the CRISPR/Cas9 method (Figure 2A & 2B). Zebrafish *tcn2* spans over 10 exons and codes for 423 amino acids long protein. The d13 line *tcn2*^-/-^ exhibited a 13 bp deletion in exon 7 (c.1087_1099 del CCAATGGCAGCAT, NM_001123231.2) leading to a frameshift mutation at amino acid 276 and a premature stop codon after 14 amino acids (Figure 2C). The d20 *tcn2*^-/-^ had a 20 bp deletion in exon 7 (c.1080_1099 del CCATAATCCAATGGCAGCAT, NM_001123231.2) leading to a frameshift mutation at amino acid 273 and a premature stop codon after 21 amino acids (Figure 2C). Immunoblotting of tcn2 during embryonic stages in wild type zebrafish revealed that it is present at all the embryonic stages (Supplementary Figure 1A). The expression levels were comparatively lower in the gastrula and 24 hpf stages. A significant decrease in *tcn2* transcripts was observed in *tcn2*^-/-^ embryos in both lines (Figure 2D). Immunoblot analysis using a polyclonal tcn2 antibody identified the ∼45 kDa band representing tcn2 in Wt embryo (Figure 2E). The band was absent in the mutant embryo lysates confirming the knockout of *tcn2*. Further, absence of the 32 kDa band in the mutant embryo lysates completely rules out any speculation of residual transcripts introduced due to deletions. Both *tcn2*^-/-^ mutant lines were phenotypically indistinguishable and hence, d13 line was used for subsequent experiments. We further checked *tcn2* expression in various adult tissues and confirmed its expression in brain, eyes, gills, muscle, heart, liver, testis, and ovary in Wt fish but complete loss of expression in the *tcn2*^-/-^ fish (Supplementary Figure 1B & 1C).

**Figure 2:**
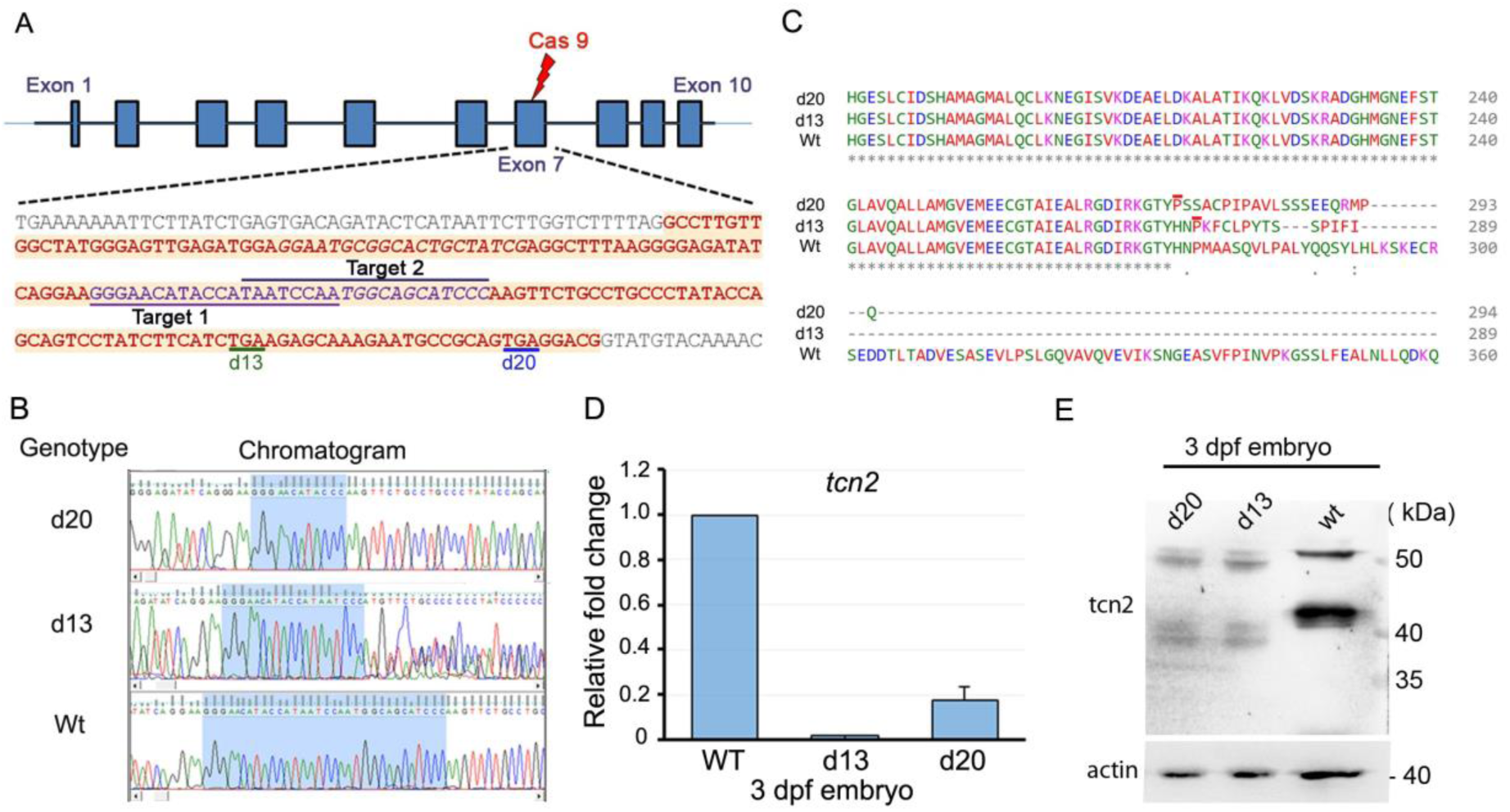
Molecular confirmation of *tcn2*^-/-^ zebrafish lines. A) Schematic showing gene structure of *tcn2* with the sequence of exon 7 (highlighted). The underlined regions are targets of cas9 gRNAs, and two premature stop codons that appeared in d13 and d20 *tcn2* mutants respectively; B) Chromatogram showing the mutated target region in the mutant embryos; C) Multiple sequence alignment showing c-termini of the predicted translated sequence of Wt and mutant *tcn2*. The red bar depicts the first affected amino acid in the mutants; D) qPCR result showing relative expression of *tcn2* transcript in 3 dpf zebrafish embryo, and, E) Immunoblot of protein lysates from wild type (Wt) and mutant embryos probed with antibodies to zebrafish tcn2 and actin.

### Discerning the *tcn2*^-/-^ phenotype and effect of B12 supplementation

The homozygous knockout embryos obtained from the cross between heterozygous parents (F1-KO) grew normally and did not show any overt phenotype. However, the progeny of the homozygous adult *tcn2*^-/-^ zebrafish (F2-KO) showed developmental deformities. Most of the F2-KO embryos showed stunted growth, remained unhatched, and were paused during the segmentation stage (Figure 3A). At 24 hpf, the F2-KO embryos were either at the early (2-8 somites) or mid-segmentation (12-16 somites) stage while the Wt embryos were at the late segmentation (28+ somites) stage. At the 2 dpf stage, the F2-KO embryos were smaller and showed developmental delay, curved bodies and tails, abnormal yolk extension, less developed caudal fin fold, and lower pigmentation. Later, the embryos developed pericardial and/or yolk sac edema. Most of the embryos were dead by 2 weeks post-fertilization stage. These results indicate that the F2-KO shows developmental defects which result in increased embryonic morbidity.

**Figure 3:**
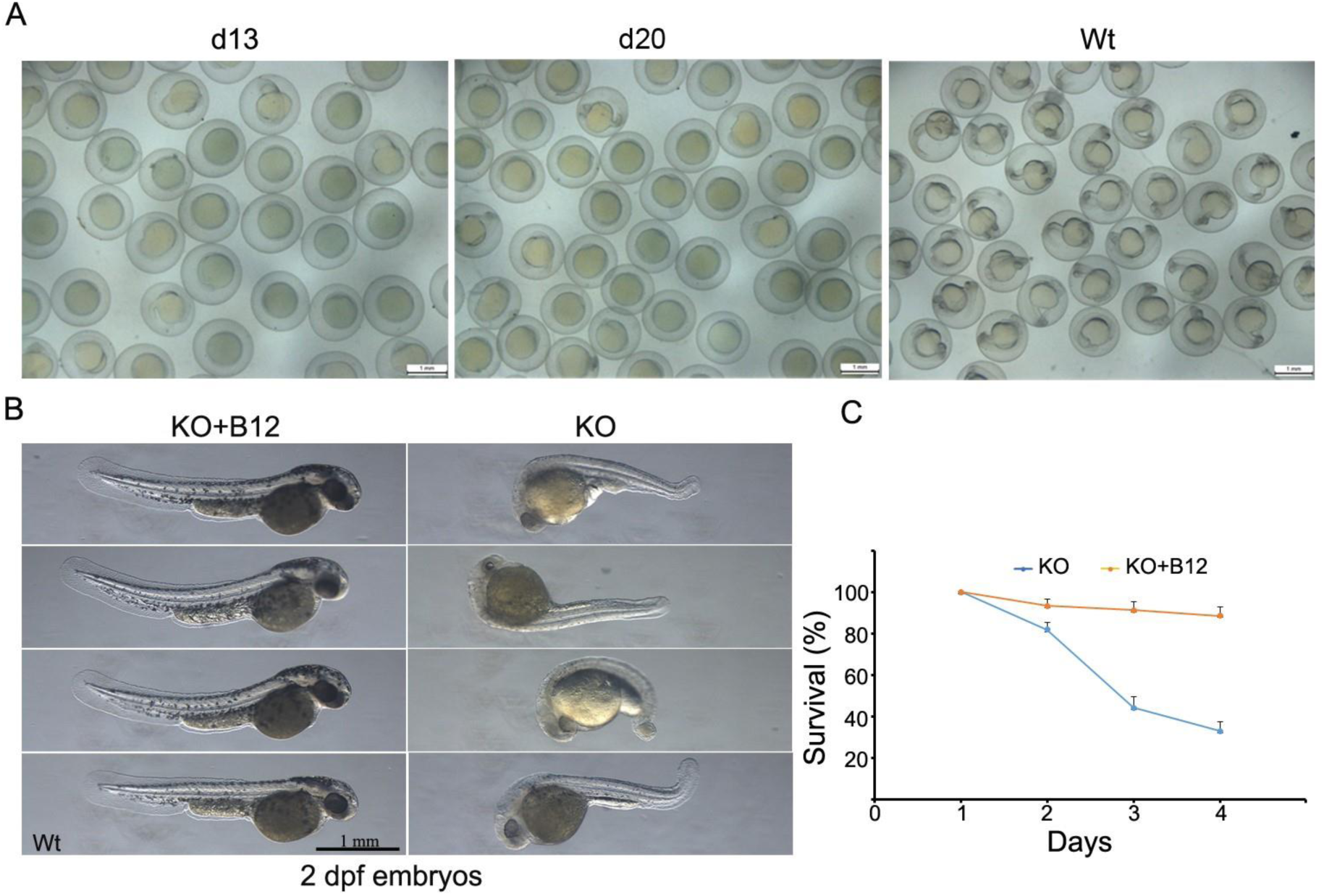
Developmental phenotypes in *tcn2*^-/-^ embryos and reversal of phenotype on B12 supplementation. (A) Developmental delay in *tcn2*^-/-^ lines at 24 hpf stage; (B) Effect of B12 supplementation on 2 dpf *tcn2*^-/-^ embryos. Images compared to a 2 dpf Wt embryo (bottom left); (C) Graph depicting the effect of B12 supplementation on the survival percentage of *tcn2*^-/-^ embryos during development. A total of 1594 embryos (700/894 for untreated/treated groups) were analysed in four batches. Error bar indicates the standard error. Majority of *tcn2*^-/-^ embryos showed stunted growth and were paused in the early segmentation stage. Bars in panel (A) & (B) indicate 1 mm.

We investigated if supplementation of B12 to the embryo medium (E3) immediately after the transfer of F2-KO embryos to the petri plates influenced the developmental phenotype observed earlier. B12 supplementation rescued the developmental defects observed in the knockout embryos (Figure 3B). Compared to the untreated embryos (∼70% death by 4 dpf stage), the B12 supplemented knockout embryos exhibited better survival rate and ∼80% of embryos were survived at 4 dpf stage (Figure 3C).

### Transcriptome analysis in *tcn2*^-/-^ reveals effect on transcription and translation processes and altered developmental and metabolic pathways

RNA-sequencing was performed to investigate the comparative transcriptomic changes occurring in Wt and the F2-KO embryos at two different stages 1-cell, and 24 hpf, with and without B12 supplementation. Across 6 samples (3 KO and 3 Wt) at 1-cell stage, at least 48.5 million reads were generated with a minimum read length of 144 bases and for 9 samples (3 Wt, 3 KO, and 3 KO+B12) at 24 hpf stage, at least 50 million reads were generated with a minimum read length of 174 bases (Supplementary Table 2B). Apart from one sample (Wt, 1-cell stage), more than 80% of these reads were successfully mapped. More than 70% of the reads were mapped uniquely for all samples except two where it was 49% and 65% (Supplementary Table S2B). We compared the RNA sequencing data from 1-cell and 24 hpf stages of *tcn2*^-/-^ embryos with their respective Wt, and also between 24 hpf stage with and without B12 to understand differentially expressed genes and associated pathways or biological processes that may be related to the developmental phenotype in the knockout embryos.

We did a technical validation of RNA-sequencing data by RT-PCR for 10-13 randomly selected DEGs representing different levels of expression based on the TPM (transcript per million) values – the *ube2m, pdp1, slc25a10b,* and *elovl7a* as representative of high expressing genes, *mtrr, acbd3,* and *sirt7* as moderate, and *cpe, pfkma* and *rbm28* as low expression genes at the 1-cell stage in zebrafish embryo [20]. Similarly, at 24 hpf stage sirt7, *pfkma, vtg, mtrr*, and *acbd3* represented as low expressing, *cpe, elovl7a, slc25a10b* and *rbm28* as moderate and *ube2m, ctsba, tpm3, hbbe3* and *apoa1b* as high expressing genes. There was a strong correlation of gene expression levels between RT-PCR and RNA-sequencing validating our RNA-sequencing results (Supplementary Figure 2).

#### Enrichment analysis of DEGs at 1-cell stage

Comparative analysis of transcriptome data from three biological samples from the 1-cell stage group of *tcn2*^-/-^ and Wt counterpart observed 3110 differentially expressed genes at log_2_FC ≥ 0.6; adjusted p-value < 0.01 of which 1434 genes were upregulated while 1676 were downregulated (Figure 4A and Supplementary Table S3A & S3B). To identify the enriched annotation terms among the DEGs, we utilized Functional Annotation Chart tool in DAVID. The GO terms or pathway enriched (FDR ≤ 0.05) in upregulated genes included pseudo-uridine synthesis, RNA modification, tRNA processing, ribosome biogenesis, spliceosome, metabolic pathways, and valine, leucine and isoleucine degradation (Table 1 and Supplementary Table S4A). For downregulated DEGs, lysosomal pathway was the most significantly enriched (Table 1 and Supplementary Table S4B).

**Figure 4:**
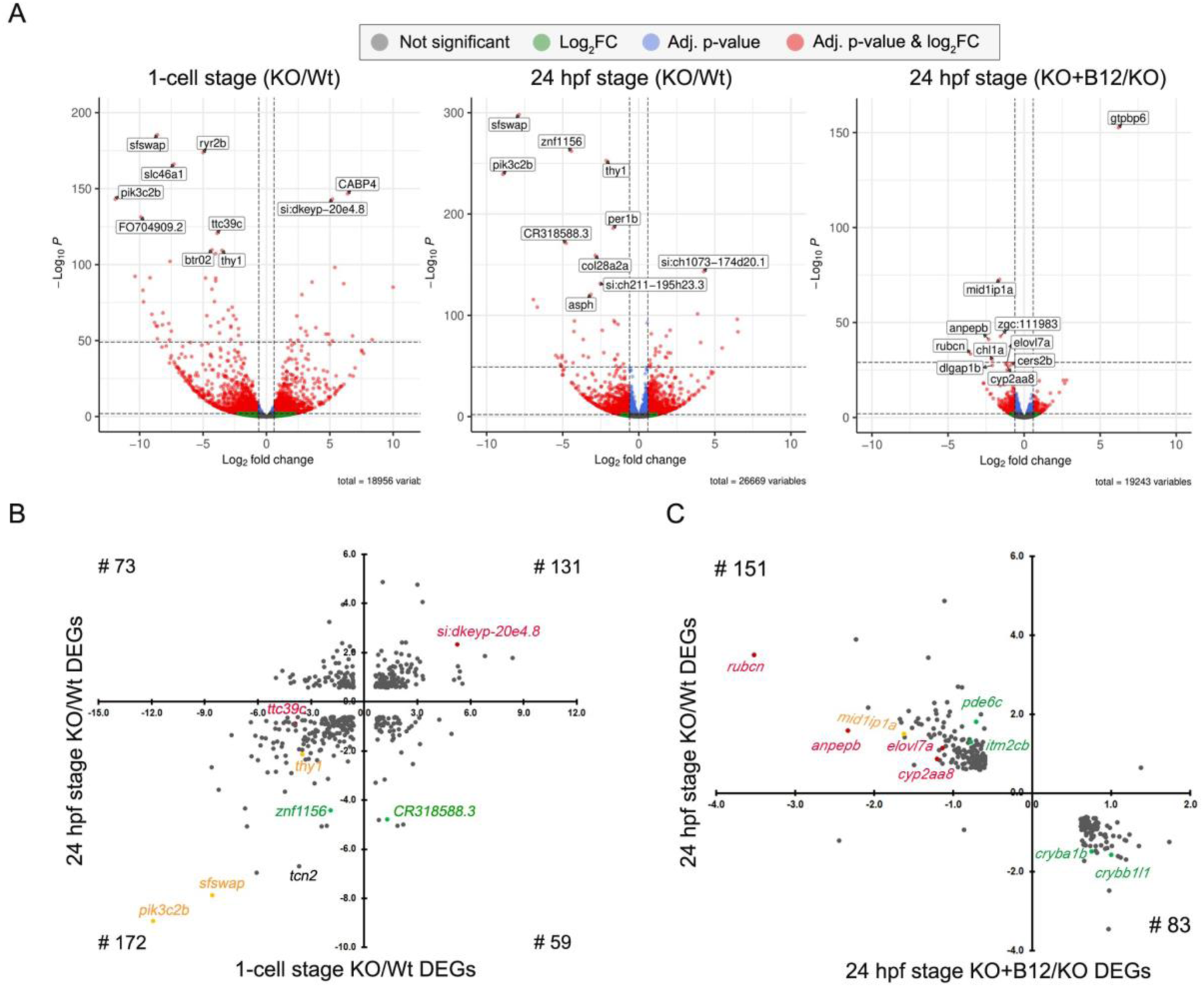
Transcriptome analysis showing differentially expressed genes at different developmental stages. (A) Volcano plots illustrating differentially express genes (DEGs) at different developmental stages. The top 10 most significant DEGs are highlighted in each group. X-axis represent log_2_FC, and y-axis represents –log10 p-value of the corresponding genes; (B) Scatter chart depicting log_2_FC values of 435 common DEGs across 1-cell and 24 hpf stage KO/Wt datasets. Quadrants I and III consist of DEGs that are upregulated (n=131) and downregulated (n=172) respectively in both stages. DEGs in Quadrant II (n=59) were upregulated in the 1-cell stage but downregulated in the 24 hpf stage while quadrant IV (n=73) were downregulated in 1- cell and upregulated in the 24 hpf stage in KO embryos; (C) A scatter chart displaying log_2_FC values for 237 common DEGs between KO/WT and KO+B12/KO at 24 hpf stage. One gene in quadrant I was upregulated and 2 genes in quadrant III were downregulated in both datasets. Genes in quadrant II (n=83) were upregulated in KO+B12/KO dataset but downregulated in KO/WT dataset and vice versa for genes in Quadrant IV (n=151).

**Table 1:** Enrichment and pathways analyses. Significantly enriched terms or pathways (FDR ≤ 0.05) from GO analysis using upregulated and downregulated genes from different groups. * The initials with ‘dre’ indicates KEGG-pathway, and ‘GO’ indicates GOTERM_BP_DIRECT.

| Stage/Group/DEG status | GO Term* | Fold Enrichment | FDR |
| --- | --- | --- | --- |
| 1-Cell (KO/Wt) upregulated | dre00280:Valine, leucine and isoleucine degradation | 3.60 | 0.027 |
|  | dre03008:Ribosome biogenesis in eukaryotes | 3.01 | 0.038 |
|  | dre01100:Metabolic pathways | 1.26 | 0.038 |
|  | dre03040:Spliceosome | 2.28 | 0.038 |
|  | GO:0001522~pseudouridine synthesis | 8.85 | 0.019 |
|  | GO:0009451~RNA modification | 9.68 | 0.019 |
|  | GO:0008033~tRNA processing | 3.54 | 0.019 |
| 1-Cell (KO/Wt) downregulated | dre04142:Lysosome | 2.13 | 0.050 |
| 24hpf (KO/Wt) Upregulated | dre01100:Metabolic pathways | 1.71 | 1.14E-04 |
|  | GO:0098869~cellular oxidant detoxification | 20.26 | 2.83E-05 |
|  | GO:0042744~hydrogen peroxide catabolic process | 15.19 | 2.83E-05 |
|  | GO:0006082~organic acid metabolic process | 7.91 | 0.016 |
|  | GO:0015671~oxygen transport | 12.80 | 0.017 |
|  | GO:0006805~xenobiotic metabolic process | 6.48 | 0.036 |
| 24hpf (KO/Wt) Downregulated | GO:0002088~lens development in camera-type eye | 8.56 | 3.2E-12 |
|  | GO:0007601~visual perception | 5.41 | 1.4E-08 |
| 24hpf (KO+B12/KO) upregulated | GO:0002088~lens development in camera-type eye | 15.39 | 5.09E-06 |
|  | GO:0007601~visual perception | 8.41 | 0.002 |
| Common between 1-cell and 24 hpf (KO/Wt) Upregulated | dre04216:Ferroptosis | 16.57 | 0.008 |
|  | dre01100:Metabolic pathways | 2.11 | 0.012 |
|  | GO:0006880~intracellular sequestering of iron ion | 91.08 | 0.002 |
|  | GO:0006826~iron ion transport | 40.98 | 0.015 |
|  | GO:0006879~cellular iron ion homeostasis | 28.26 | 0.032 |
| Downregulated in KO/Wt but upregulated in KO+B12/KO | GO:0002088~lens development in camera-type eye | 25.73 | 2.83E-08 |
|  | GO:0007601~visual perception | 12.50 | 2.99E-04 |

Developmental processes are greatly influenced by complex interactions between intrinsic factors i.e., genetics and epigenetics, and extrinsic factors like maternal nutrition. We examined the dysregulated epigenetic regulators among the DEGs at 1-cell stage and identified 85 genes (comprising 43 upregulated and 42 downregulated genes; Supplementary Table 5) that had a human ortholog known to have an epigenetic function in the EpiFactor database [21]. These genes included chromatin remodelers, histone modification writers/erasers/readers, RNA modifiers, etc. Among chromatin remodelers, *ino80e, ctcf, bptf, ss18l2, ercc6, actr3b* etc., were upregulated and *rai1, cul2, hmg20b, ctbp1, chd9, ncoa2,* etc were downregulated (Supplementary Table 5). Notably, a group of genes involved in RNA modifications like *mettl3, trub2, rsrc1, alkbh5, exosc6,* and *rbm11* were upregulated whereas *sfswap, nsun6, celf4,* and *mbnl3* were downregulated (Supplementary Table 5).

#### Enrichment analysis of DEGs at 24 hpf stage

An analysis of RNA-Seq data on three biological samples from *tcn2*^-/-^ and Wt embryos at the 24 hpf stage identified 2741 annotated DEGs (Figure 4A and Supplementary Table S3A & S3C). Of these, 1261 were upregulated and 1480 were downregulated, with 610 and 728 genes respectively mapped by DAVID. The upregulated genes were significantly enriched in metabolic pathways, cellular oxidant detoxification, organic acid metabolic process, oxygen transport, and xenobiotic metabolic process (Table 1 and Supplementary Table 4C). For downregulated genes, lens development in camera-type eye, and visual perception were the most enriched biological processes (Table 1 and Supplementary Table 4D).

#### Comparative analysis of DEGs at 1-cell and 24 hpf stages

In *tcn2*^-/-^ embryos across 1-cell and 24 hpf stages, there were 435 common differentially expressed genes. Three genes – *sfswap* (log_2_FC = −8.6 & −7.9, p-value = 4.2e-186 & 9.0e-299), *thy1* (log_2_FC = −3.5 & −2.1, p-value = 4.9e-110 & 2.0e-253) and *pik3c2b* (log_2_FC = −11.92 & −8.93, p-value = 8.2e-144 & 6.4e-240) were common in top ten most significant genes in both 1-cell and 24 hpf (Figure 4A and Supplementary Table S3E). Among these 435 common DEGs, 303 genes demonstrated a congruent directionality in their expression changes with 131 being upregulated and 172 down-regulated (Figure 4B). Additionally, a subset of 132 genes exhibited an inversely regulated pattern of expression. The common upregulated genes during 1-cell and 24 hpf were significantly enriched in metabolic pathways and iron metabolism including ferroptosis, intracellular sequestering of iron ion, iron ion transport and cellular iron homeostasis (Table 1 and Supplementary Table S4E). No significantly enriched pathways were identified for commonly downregulated genes in 1-cell and 24 hpf stages (Supplementary Table S4F).

#### Influence of B12 supplementation on DEGs in tcn2^-/-^ embryos at 24 hpf

The transcriptome of *tcn2*^-/-^ embryos was compared to that of *tcn2*^-/-^ embryos supplemented with B12 at the 24 hpf stage (Figure 4A). The analysis identified 422 DEGs, of which 237 were common to those identified in the 24 hpf KO/Wt dataset (Figure 4C and Supplementary Table S3A & S3F). Except for the three DEGs, all the other 234 genes revealed restoration of gene expression akin to wild type levels suggesting a phenomenon of transcriptional reversion facilitated on B12 supplementation (Figure 4C and Supplementary Table S3F). Functional annotation and enrichment analyses of those 83 genes which were upregulated in B12 supplementation but downregulated in non-supplementation were enriched in lens development in camera-type eye, and visual perception as the most significant biological processes (Table 1 and Supplementary Table S4G). We did not observe significant enrichment for those 151 genes which were downregulated in B12 supplementation compared to non-supplementation group. Although they did not pass the FDR, many other top pathways were also reversed after B12 supplementation included synaptic transmission, glycinergic, negative regulation of DNA recombination, chromosome condensation, metabolic pathways, signaling pathways, biosynthesis of unsaturated fatty acids, and fatty acid elongation (Supplementary Table S4H).

## Discussion

In this study, we demonstrate the critical role of B12 in early developmental processes by generating *tcn2* loss of function alleles in zebrafish by the CRISPR/cas9 method. Both mutant lines showed significant reductions in *tcn2* transcript which was validated by immunoblot analysis confirming a successful knockout of *tcn2*. We assessed the phenotypic and transcriptomic changes during embryonic development in normal and B12 deficient states, and investigated the effect of B12 supplementation on the reversal of phenotype and underlying transcriptomic changes.

We have made several interesting observations. The first-generation *tcn2*^-/-^ zebrafish (F1-KO) did not show any abnormal phenotype. However, when these *tcn2*^-/-^ fish were allowed to breed, the embryos (F2-KO) showed delayed development with a range of deformities like curved tail, pericardial and yolk sac edema, etc., and increased mortality; similar to the observation reported by Benoit et al. [22,23]. Majority of F2-KO embryos were developmentally stalled at early or mid-segmentation stage, and nearly 70% larvae were dead by completion of day 4, with a sharp increase in deaths on day 6 as reported earlier [22]. The variations in survival rates between the two studies cannot be attributed to a single clear explanation. Nevertheless, these differences may arise from variances in the knockout target site within *tcn2*, as well as variations in the rearing conditions and the type of feed provided to the fish. The immunoblot analysis conducted at various stages of embryonic development revealed that *tcn2* is a maternally loaded gene as evidenced by its presence at 1-cell stage. The presence of tcn2 at all the embryonic developmental stages suggests that zygotic *tcn2* expression may commence as early as maternal zygotic transitions. It may not be unreasonable to surmise that maternally loaded *tcn2* mRNA or tcn2 protein in the egg might have rescued *tcn2*^-/-^ phenotype in the F1-KO embryos, and its unavailability in the F2-KO embryos may have resulted in morbidity and early mortality of these embryos. This was further supported by Benoit et al. that the developmental defect in embryos from the knockout female were independent of male genotype [22]. Further, the rescuing of developmental defect in F2-KO by B12 supplementation suggests a direct effect of B12 in early embryonic development. We hypothesize that maternal tcn2 is important for the reserve of B12 in the eggs that is essential for early embryonic development. However, normal development of F1-KO and their attainment of sexual maturity suggests that there could be compensatory mechanisms in *tcn2*^-/-^ zebrafish at a later stage. This needs further investigation.

Interestingly, F2-KO embryos developed normally until gastrulation stage but were stalled during early or mid-segmentation stage and thus affecting the differentiation and organogenesis [24]. To gain insights into the molecular mechanisms underlying these phenotypic disparities, we performed transcriptome analysis at two developmental stages. Transcriptome analysis at 1-cell stage showed differential gene expression in the maternally loaded transcripts. Previously, single-cell RNA-seq analysis has revealed that mature oocytes in zebrafish were enriched in the ribosome, spliceosome, oxidative phosphorylation, RNA transport, oocyte meiosis, pyrimidine metabolism, etc. [25]. This emphasizes the importance of these pathways during early embryonic development. Interestingly we observed that genes involved in various transcription and translation related processes such as RNA modification, tRNA processing, spliceosome, ribosome biogenesis, and amino acid catabolism etc., were upregulated in *tcn2*^-/-^ embryos indicating impact on protein synthesis and energy homeostasis etc., and that may have detrimental cascading effect on cell proliferation. This observation gains further support from the observed downregulation of histone deacetylase genes (*hdac5, hdac9b and hdac8*) observed in knockout embryos at 1-cell stage and downregulation of HDACs associated with decreased cell proliferation [26]. These observations were consistent with the reduction in cell proliferation in neuroblastoma cells [27] and impaired DNA synthesis in erythroblasts [28] in B12 deficiency. The downregulated genes were significantly enriched in lysosome pathways and interestingly lysosomes play a unique role in nutrient mobilization, cellular differentiation, and maintenance of homeostasis and signaling during embryonic development [29,30].

The limited supply of maternally deposited transcripts during transcriptionally silent developmental stages suggests that post-transcriptional control mechanisms may play a crucial role in translating these transcripts in response to dynamic changes during early embryonic divisions [31–33]. In the knockout embryos, a global dysregulation was observed in genes involved in epigenetic regulation including chromatin remodeling, histone post-translational modifications, histone chaperones, and DNA and RNA modifications. Many of these genes were moderately or highly expressed in the 1-cell stage and a small perturbation in the expression could affect early embryonic development. Among the top 10 most significant DEGs, *sfswap* (splicing factor SWAP), *thy1* (Thy-1 cell surface antigen) and *pik3c2b* (phosphatidylinositol 4-phosphate 3-kinase C2 domain-containing subunit beta) status was shared in both 1-cell and 24 hpf stages. The *sfswap* encodes an RS-domain containing (SR-Like) protein that regulates RNA processing including splicing, transcriptional elongation, genome and transcript stability, miRNA processing, mRNA degradation, and translational controls, etc. [34]. It is further known to affect Notch signaling pathway and inner ear development in mice [35]. *Thy1* encodes a cell surface glycoprotein and member of the immunoglobulin superfamily of proteins which is involved in cell adhesion and communication in the immune and nervous systems. Previous studies have shown that *Thy1* is an m^6^A-modified gene and has important roles in neuronal differentiation and morphology [36,37]. *Pik3c2b* encodes a protein belong to class II phosphoinositide 3-kinase (PI3K) subfamily, and plays a role in vesicle trafficking cell signaling, cell division, proliferation, and survival etc.[38]. It was interesting to observe that the key epigenetic regulatory genes, such as *ctcf*, which encodes the zinc finger binding protein ctcf that is involved in chromosome folding and insulation of topologically associated domains was distinctly upregulated (log_2_FC=4.3) in 1-cell *tcn2*^-/-^ embryos. CTCF is also involved in the regulation of miRNAs, plays a crucial role in the pluripotency of cells, and is linked to the modulation of alternative splicing at the transcriptome-wide level [39,40]. Two maternal genes, *mettl3* (encodes N6-adenosine-methyltransferase complex catalytic subunit) and *alkbh5* (encodes alkB homolog 5, RNA demethylase) were moderately but significantly upregulated (log_2_FC = 0.61 and 0.81 respectively) in *tcn2*^-/-^ embryos. These genes play a crucial role by oppositely catalyzing m^6^A RNA methylation which affects the stability of the modified mRNAs, splicing, nuclear export, mRNA translation and are critical in fate determination during embryogenesis [41,42].

At 24 hpf, the maternally loaded transcripts were depleted and thus transcripts at this stage mostly represented the zygotic transcripts. Genes involved in oxidative pathways including cellular oxidant detoxification, oxygen transport and H_2_O_2_ catabolic pathways were upregulated whereas eye development, visual perception, and synaptic transmission associated genes were down regulated and correlated with the observed phenotypes. These observations were similar to the reported findings in 2 dpf *tcn2*^-/-^ zebrafish transcriptome analysis [22]. Interestingly, many of the DEGs at 1-cell stage continued to exhibit differential expression at 24 hpf signifying a persistent pattern of DEG alterations across both maternally loaded and zygotic transcripts. The common upregulated DEGs at two embryonic stages, 1-cell and 24 hpf were enriched in metabolic pathways and iron metabolism such as ferroptosis, cellular iron transport and iron sequestration. Increased ferroptosis leads to oxidative stress which can potentially disrupt essential cellular functions during embryo development. These findings were consistent with a recent study in *C. elegans* that increased lipogenesis and peroxidation in adult worms, leading to germline defects through ferroptosis due to early life B12 deficiency [43]. The common downregulated DEGs were enriched in visual perception, and lens development as it was reported in [22] and correlated with the phenotypic expression. Transcriptome analysis in B12 supplemented knockout embryos showed the reversal effect of many of DEGs observed in 24 hpf knockout embryos indicating a direct or indirect effect of B12 supplementation on expression of these genes and their probable role in rescuing the developmental defects. For example, the visual perception and lens development pathways genes were upregulated in B12 supplementation knockout compared to non-supplemented knockout groups at 24 hpf stage. This finding indicates the role of B12 in early embryonic development by possibly regulating these differentially expressed genes. The rescuing of phenotype and reversal of gene expressions by B12 supplementation revealed that B12 plays an important role in epigenetic regulation including RNA processing and stability thereby maintaining transcription and translation during early embryonic development. It will be interesting to investigate whether these DEGs are the cause or effect of the developmental delays in knockout embryos. B12 is a key metabolite in OCM which maintains the methylation potential in cell. Our previous studies of B12 intervention in children and preconception mothers had shown the differential DNA methylation in genes associated with lipid metabolism, type 2 diabetes, and its intermediate traits [44,45]. Therefore, it will be worth investigating DNA and RNA methylation changes in these DEGs during the embryonic developmental stages to understand the underlying molecular regulation.

### Strengths and limitations

The complexity of the knockout phenotypes makes it challenging to draw precise conclusions regarding the effect of *tcn2* knockout on the transcriptome and causality of developmental delay. The biggest strength of this study is understanding the molecular changes at two different stages of embryonic development in B12 deficiency models and further, restoration of molecular changes and the phenotypes thus making one of the most robust contributions in the role of B12 in early embryonic development. Replication of some of the transcriptomic changes from earlier studies make the conclusion stronger. Certainly, a comprehensive metabolic profile including B12 levels and other related metabolites in eggs or embryos, and adult zebrafishes would have strengthened our conclusions and provided a better insight into the effect of *tcn2* knockout at metabolite levels. We do not know what compensatory mechanisms rescue the F1-KO embryos as well as the knockout adult fishes without B12 supplementation, and this need further investigation. Since the B12 supplementation was provided throughout the embryonic developmental stages, hence it is difficult to pinpoint the critical stage where the supplementation would have the best effect. Functional validation of targeted DEGs will be required to elucidate the role of B12 in transcriptional and translational regulations. Analysis on epigenetic markers including DNA and RNA methylation during the embryonic developmental stages will provide a comprehensive picture of transcriptional regulation.

### Conclusions

Through a *tcn2*^-/-^ zebrafish model for B12 deficiency, we have provided comprehensive molecular evidence on the essentiality of vitamin B12 during early embryonic development. The *tcn2* knockout leads to altered gene expressions involved in epigenetic regulation, transcription and translation machineries, and various metabolic pathways which are critical during embryonic development. The reversal effect at both phenotypic as well as the gene expression levels by B12 supplementation highlights the importance of B12 in early developmental processes. This study provides novel insights into the gene dysregulation of the maternal and zygotic transcriptome in *tcn2*^-/-^ embryos which will add to our understanding of the role of maternal B12 levels during pre and early pregnancy on the development of the embryo. This is important in context of humans as a large percentage of women at childbearing age are B12 deficient, and thus an early intervention will be essential for a healthy fetal development.

## Acknowledgements

We thank Council of Scientific and Industrial Research, Department of Scientific and Industrial Research (DSIR), Ministry of Science and Technology, Govt of India, New Delhi for financial support to ADV, SSN, MC, AU and SR. GRC conveys his sincere gratitude to Science and Engineering Research Board (SERB), Department of Science and Technology (DST), Ministry of Science and Technology, Govt of India, New Delhi for the award of JC Bose Fellowship (JCB/2021/000042).

## Contributions

The study was conceived by GRC, ADV and SSN. ADV and SSN designed the strategy for generating the zebrafish knock out embryos and their further characterization. ADV conducted all experiments on zebrafish with help from MC and SR. AU contributed to analysis of the transcriptome data on the embryos. ADV wrote the first draft with significant inputs from SSN and GRC. The manuscript has been seen and approved by all the authors. GRC is the guarantor of the study.

---

**Supplementary Table 1:** Oligonucleotides used in the study.

**Supplementary Table 2:** RNA sequencing quality control parameters and summary of sequencing read statistics.

**Supplementary Table 3:** Differentially expressed genes obtained in the RNA sequencing analysis at different stages.

**Supplementary Table 4:** Functional annotation and enrichment analysis of differentially expressed genes from different developmental stages.

**Supplementary Table 5:** Differentially expressed genes from 1-cell stage with human orthologs known to have role in epigenetic regulation.

**Supplementary Figure 1:**
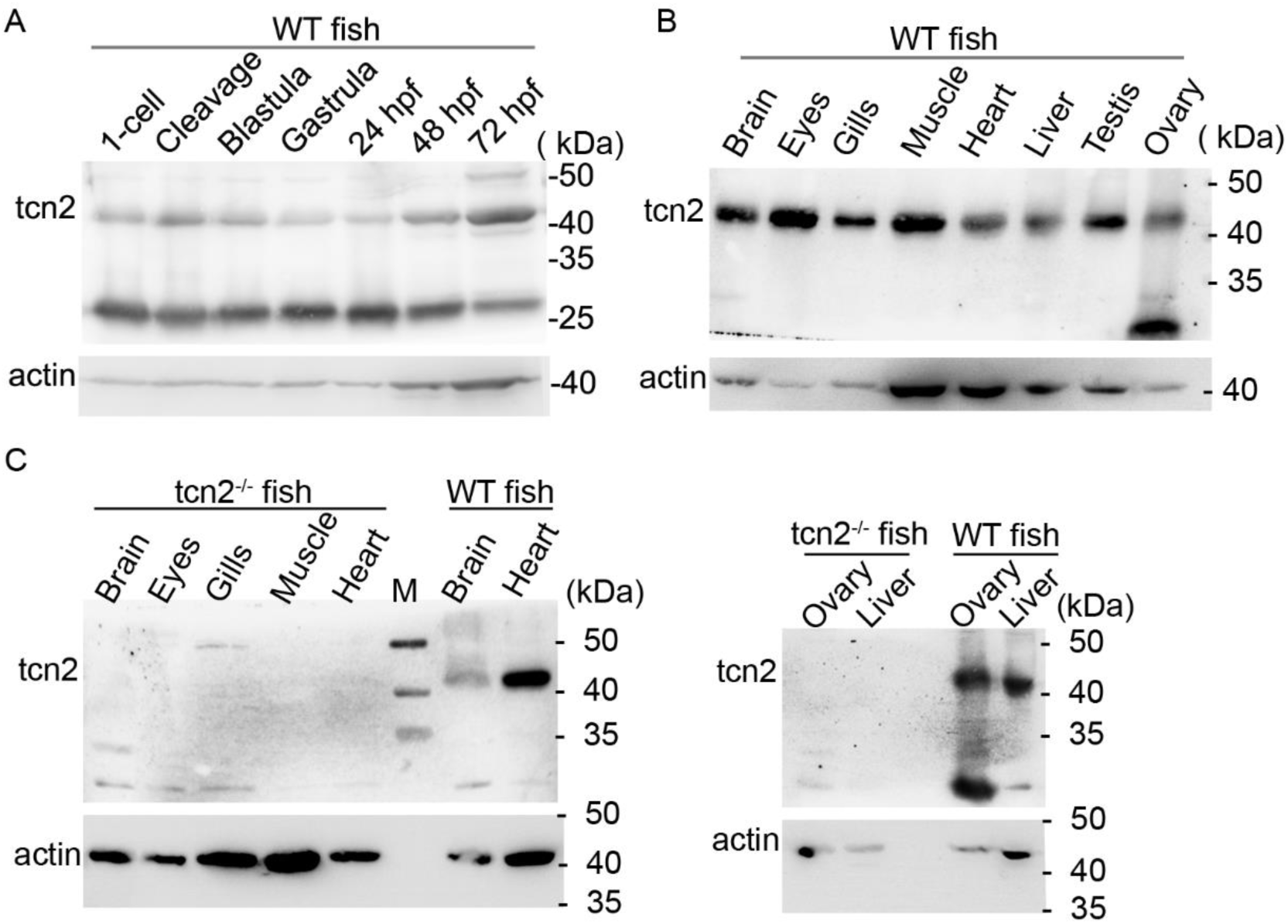
Embryonic and tissue-wide expression of tcn2 in zebrafish. A) Immunoblot of lysates of different embryonic stages (1-cell to 72 hpf stage) probed with antibody against actin and in-house raised polyclonal antibody against zebrafish tcn2. Apart from the specific band at ∼43kDa, a band is also observed at ∼30 kDa. B) Immunoblot showing expression of tcn2 in different tissues, and, C) Immunoblot showing comparative expression of tcn2 in different tissues in tcn2^-/-^ and wild type zebrafish.

**Supplementary Figure 2:**
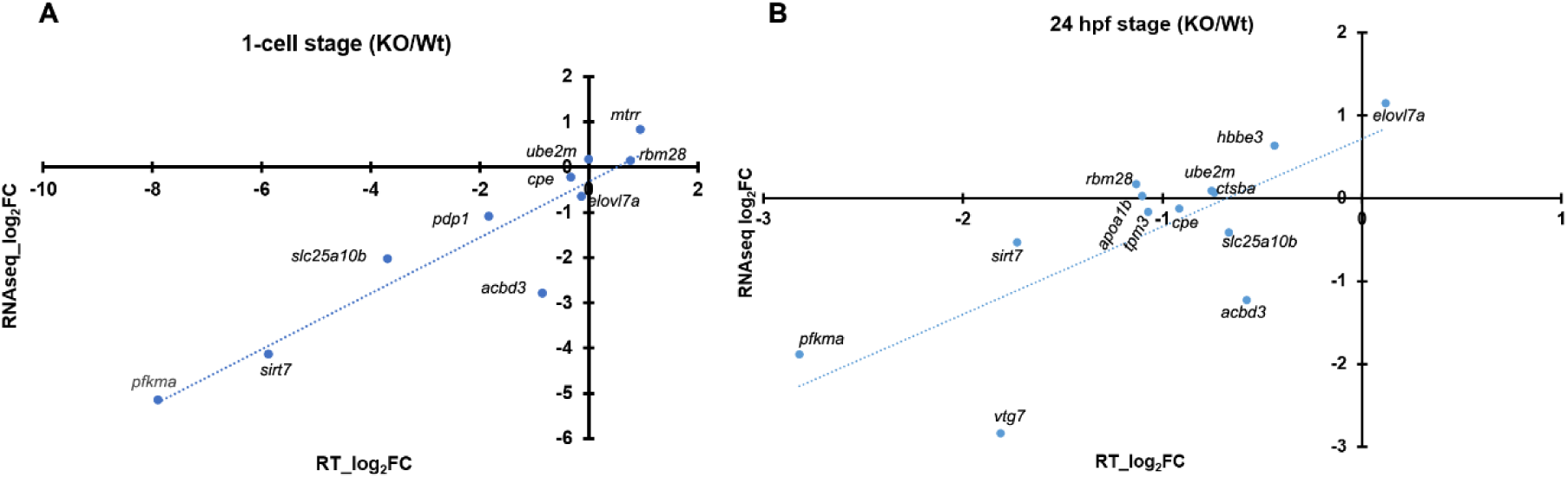
Comparative log_2_fold change value of genes between RNAseq and Real time analysis. The scatter plot depicts log_2_ fold change values of different genes across two analyses. *pfkma*, phosphofructokinase, muscle a; *sirt7*, sirtuin7; *slc25a10*, solute carrier family 25 member 10b; *mtrr*, 5-methyltetrahydrofolate-homocysteine methyltransferase reductase; *cpe*, carboxypeptidase E; *pdp1*, pyruvate dehydrogenase phosphatase catalytic subunit 1; *acbd3*, acyl-Coenzyme A binding domain containing 3; *evovl7a*, ELOVL fatty acid elongase 7a; *ube2m*, ubiquitin conjugating enzyme E2 M; *rbm28*, RNA binding motif protein 28; *vtg*, vitellogenin; *tpm3*, tropomyosin 3; *apoa1b*, apolipoprotein A-lb; *ctsb*, cathepsin Ba, *hbbe3*, hemoglobin beta embryonic-3.

